# Antibody-mediated VWF targeting reconstitutes ADAMTS13 metalloproteinase activity and prevents thrombocytopenia in mice

**DOI:** 10.64898/2026.09.11.750821

**Authors:** Atsushi Hashimoto, Yoshiki Komeda, Fumika Kiku, Minako Oogi, Mariko Nakano, Sagano Shiina, Harue Imai, Yasuhisa Shiraishi, Kazuhiro Masuda, Akiko Sano

## Abstract

ADAMTS13 (a multidomain metalloproteinase) cleaves von Willebrand factor (VWF), thereby preventing the accumulation of ultra-large VWF multimers that promote platelet-rich thrombus formation. Severe ADAMTS13 deficiency causes thrombotic thrombocytopenic purpura (TTP). In immune-mediated TTP (iTTP), anti-ADAMTS13 autoantibodies inhibit ADAMTS13 activity, often by targeting non-catalytic domains that mediate substrate recognition. We developed ADAMTS13 mimetics by fusing the ADAMTS13 metalloproteinase domain to anti-human VWF A2 domain antibodies, aiming to replace native substrate-recognition domains with antibody-mediated substrate targeting while removing major autoantibody-targeted ADAMTS13 domains. The resulting mimetics bound to the recombinant VWF A2 domain and showed proteolytic activity toward both the recombinant VWF A2 domain and FRETS-VWF73 substrate. Kinetic analysis showed that the mimetics had not only lower *k*_cat_ values than ADAMTS13 but also lower *K*_m_ values, thereby maintaining overall catalytic efficiency. Importantly, ADAMTS13 mimetics retained VWF A2-cleaving activity in the presence of a spacer domain-directed anti-ADAMTS13 autoantibody both *in vitro* and *in vivo*. In an iTTP mouse model, ADAMTS13 mimetics prevented recombinant VWF-induced thrombocytopenia, whereas the parental anti-VWF A2 domain antibody did not. Platelet-preserving activity was also observed in ADAMTS13-deficient mice. In further analyses, some mimetics showed reduced binding to a conformationally restricted VWF A2 domain, resulting in reduced cleavage activity. Antibody binding correlated with platelet recovery in the iTTP mouse model, suggesting that epitope accessibility and antibody-mediated positioning contribute to *in vivo* activity. These findings demonstrate that antibody-mediated substrate targeting can reconstitute the VWF-cleaving activity of the ADAMTS13 metalloproteinase domain and suggest that ADAMTS13 mimetics represent a potential therapeutic approach for iTTP.

## Introduction

ADAMTS13 (A disintegrin and metalloproteinase with a thrombospondin type 1 motif 13) is a multidomain metalloproteinase that regulates the multimeric size and function of von Willebrand factor (VWF), a glycoprotein essential for primary hemostasis and platelet adhesion.^1-4^ ADAMTS13 consists of a metalloproteinase domain, a disintegrin-like domain, a cysteine-rich domain, a spacer domain, eight thrombospondin type 1 repeats, and two CUB (Complement C1r/C1s, Uegf, and Bone morphogenetic protein 1) domains.^1-3^ The disintegrin-like, cysteine-rich, and spacer domains are required for efficient substrate recognition, and mutations in these domains reduce catalytic efficiency.^5-12^ Under static conditions, VWF remains in a compact conformation, which limits access of ADAMTS13 to its cleavage site.^13,14^ Under high shear stress, VWF unfolds and exposes the Tyr1605-Met1606 scissile bond in the A2 domain, enabling cleavage by ADAMTS13.^13,14^

Severe ADAMTS13 deficiency causes thrombotic thrombocytopenic purpura (TTP), a rare but life-threatening disease characterized by thrombocytopenia and microangiopathic hemolytic anemia.^15-17^ ADAMTS13 deficiency leads to the accumulation of ultra-large VWF multimers and subsequent formation of platelet-rich microvascular thrombi.^17^ Approximately 95% of TTP cases are immune-mediated TTP (iTTP), caused by anti-ADAMTS13 autoantibodies, whereas 5% are congenital TTP, caused by pathogenic variants in the *ADAMTS13* gene.^16,18-20^ Recombinant ADAMTS13 has been established as an enzyme-replacement therapy for congenital TTP.^21^ In iTTP, however, restoring ADAMTS13 activity remains challenging because anti-ADAMTS13 autoantibodies can inhibit full-length ADAMTS13. The current standard treatment for iTTP includes daily therapeutic plasma exchange, immunosuppressive therapy, and caplacizumab, an anti-VWF A1 domain nanobody.^22,23^ Although these therapies have markedly improved clinical outcomes, current treatment strategies do not directly restore ADAMTS13 proteolytic activity in the presence of inhibitory autoantibodies, highlighting the need for alternative approaches that can reconstitute the VWF-cleaving activity.^22,23^

Epitope-mapping studies have shown that anti-ADAMTS13 autoantibodies in patients with iTTP frequently target the spacer domain.^24-27^ In particular, a cluster of surface-exposed residues within the spacer domain, including Arg568, Phe592, Arg660, Tyr661, and Tyr665, forms a key exosite for interaction with the VWF A2 domain and is frequently recognized by pathogenic autoantibodies.^28-31^ These findings provided a rationale for engineering ADAMTS13 variants with spacer-domain mutations that reduce autoantibody binding.^28,30^ However, comprehensive mutational analysis of these spacer-domain residues revealed a trade-off between enzymatic activity and escape from autoantibody recognition.^31^ Thus, developing ADAMTS13 variants that evade anti-spacer-domain autoantibodies while retaining full enzymatic activity remains challenging.

In this study, we sought to develop fusion proteins, hereafter referred to as ADAMTS13 mimetics, by fusing the ADAMTS13 metalloproteinase domain to a VWF-targeting antibody. Because the disintegrin-like, cysteine-rich, and spacer domains primarily mediate VWF substrate recognition, we hypothesized that a VWF-targeting antibody could functionally replace these non-catalytic domains and thereby reconstitute the VWF-cleaving activity. We show that ADAMTS13 mimetics cleave VWF substrates *in vitro* and prevent thrombocytopenia in a mouse model of TTP. These findings suggest that ADAMTS13 mimetics may provide a potential therapeutic approach for iTTP.

## Methods

### Protein expression and purification

Synthetic DNA fragments encoding ADAMTS13 mimetics were cloned into an in-house mammalian expression vector derived from the pCI Mammalian Expression Vector (Promega, WI, USA). Expression plasmids were transiently transfected into Expi293F cells (Thermo Fisher Scientific, MA, USA), and the cells were cultured at 37°C in a humidified atmosphere containing 5% or 8% CO_2_. Culture supernatants were harvested 3-4 days after transfection and clarified by centrifugation. ADAMTS13 mimetics were purified from clarified culture supernatants using MabSelect SuRe resin (GE Healthcare, IL, USA). Purified proteins were buffer-exchanged into acetate formulation buffer consisting of 50 mM sodium acetate, 250 mM *D*-sorbitol, 50 mM *L*- arginine, and 44 μM ZnSO_4_ (pH 5.0) using NAP columns (GE Healthcare, IL, USA).

Recombinant human VWF A2 domain proteins were purified from culture supernatants using cOmplete His-Tag Purification resin (Roche, Basel, Switzerland) (Supplemental Table 1). Bound proteins were eluted with 250 mM imidazole-containing buffer and subsequently buffer-exchanged into phosphate-buffered saline (PBS). Protein size and purity were assessed by sodium dodecyl sulfate-polyacrylamide gel electrophoresis (SDS-PAGE).

### SPR analysis of ADAMTS13 mimetic and parental antibody binding to VWF A2

Binding kinetics of ADAMTS13 mimetics and parental anti-VWF A2 antibodies to recombinant VWF A2 were analyzed by surface plasmon resonance (SPR) using HBS-EP+ buffer (Cytiva, Marlborough, MA, USA) as the running buffer. An anti-human fragment crystallizable (Fc) antibody was immobilized onto a CM5 sensor chip (Cytiva) by amine coupling using a Human Antibody Capture Kit (Cytiva). Antibodies or ADAMTS13 mimetics were captured on the immobilized anti-human Fc antibody surface. A flow cell without captured antibody or ADAMTS13 mimetic was used as the reference surface. Recombinant VWF A2 was injected as the analyte at a flow rate of 30 µL/min. VWF A2 was prepared by 2.5-fold serial dilution with a top concentration of 250 nM. Association and dissociation phases were monitored for 120 s and 300 s, respectively. Binding kinetics were determined using a single-cycle kinetics method, and sensorgrams were fitted using a 1:1 binding model.

### Binding assays with VWF A2 domain

Binding of ADAMTS13 mimetics to recombinant VWF A2 domain proteins was evaluated by an enzyme-linked immunosorbent assay (ELISA). Recombinant human VWF A2 domain proteins were diluted to 2 µg/mL in PBS and immobilized on Nunc MaxiSorp 96-well plates (Thermo Fisher Scientific, MA, USA) overnight at 4°C. The plates were washed with PBS containing 0.05% Tween 20 (PBS-T) and blocked with Blocking Buffer (1% bovine serum albumin [BSA] and 2% fetal bovine serum [FBS] in PBS) for 2 h at room temperature. After washing with PBS-T, serial dilutions of ADAMTS13 mimetics prepared in Blocking Buffer were added to the plates and incubated for 2 h at room temperature. The plates were then washed with PBS-T and incubated with a horseradish peroxidase (HRP)-conjugated goat anti-human immunoglobulin G (IgG) Fc-specific antibody (IBL, Gunma, Japan) diluted 1:4000 in Blocking Buffer for 1 h at room temperature. After washing with PBS-T, bound ADAMTS13 mimetics were detected using a TMB (3,3′,5,5′-tetramethylbenzidine) substrate (Dako, CA, USA), and the reaction was stopped with 0.5 M H_2_SO_4_ (FUJIFILM Wako Pure Chemical Corporation, Osaka, Japan). Absorbance was measured at 450 nm with background correction at 570 nm.

### ADAMTS13 activity assays with recombinant VWF A2 domain

ADAMTS13 activity against the recombinant VWF A2 domain was evaluated using ELISA-based cleavage assays. Nunc MaxiSorp 96-well plates were coated with an anti-His antibody (R&D Systems, Minneapolis, MN, USA) or anti-red fluorescent protein (RFP) antibody (Rockland Immunochemicals, Pottstown, PA, USA) at 5 µg/mL in PBS for 2 h at room temperature, washed with PBS-T, and blocked with Blocking Buffer for 2 h at room temperature. The N-terminal His_8_- tagged recombinant human VWF A2 domain or mCherry-tagged recombinant human VWF A2 domain was diluted to 2 µg/mL in Blocking Buffer and immobilized on the corresponding antibody-coated plates overnight at 4°C. After washing, recombinant ADAMTS13 (R&D Systems), ADAMTS13 mimetics, the ADAMTS13 metalloproteinase domain alone, or parental anti-VWF A2 antibodies were added in FRET buffer (5 mM bis(2-hydroxyethyl)iminotris(hydroxymethyl)methane [Bis-Tris], 150 mM NaCl, 25 mM CaCl_2_, 0.005% Tween-20, pH 7.2) and incubated for 3-5 h at room temperature. Cleavage of the VWF A2 domain was detected using an HRP-labeled anti-human VWF A2 ADAMTS13-cleaved antibody (R&D Systems). The antibody was labeled with HRP using a Peroxidase Labeling Kit-NH_2_ (Dojindo Laboratories, Kumamoto, Japan) according to the manufacturer’s instructions and diluted 1:4000 in Blocking Buffer. After incubation for 1 h at room temperature, plates were washed with PBS-T, and TMB substrate (Dako, Carpinteria, CA, USA) was added. The reaction was stopped with 0.5 M H_2_SO_4_ (FUJIFILM Wako Pure Chemical Corporation, Osaka, Japan), and absorbance was measured at 450 nm with background correction at 570 nm.

### Fluorogenic ADAMTS13 activity assay

Enzyme-specific activity was determined using the FRETS-VWF73 peptide substrate (PEPTIDE INSTITUTE, INC., Osaka, Japan) in a fluorogenic assay. FRETS-VWF73 and test samples were diluted in FRET buffer (5 mM Bis-Tris, 150 mM NaCl, 25 mM CaCl_2_, 0.005% Tween-20, pH 7.2), mixed 1:1 (50 µL + 50 µL) in microplate wells, and incubated at room temperature. FRETS-VWF73 was diluted in FRET buffer to prepare a two-fold serial dilution from 32 µM to 0.25 µM. ADAMTS13 mimetics and recombinant human ADAMTS13 were adjusted to 200 nM and 60 nM, respectively. Reaction mixtures were mixed on a plate shaker, and fluorescence (excitation 340 nm, emission 450 nm) was recorded every 5 min for 60 min using a microplate reader. To convert fluorescence into the absolute amount of cleavage product, a calibration curve was generated using equimolar mixtures of FRETS-25-STD1 (PEPTIDE INSTITUTE, INC.) and FRETS-25-STD2 (PEPTIDE INSTITUTE, INC.). The initial velocity was calculated for each substrate concentration and used for kinetic analysis. Michaelis-Menten kinetic parameters were estimated using JMP Pro software version 16-17 (SAS Institute, Cary, NC, USA).

### iTTP mouse model

All animal experiments were approved by the Institutional Animal Care and Use Committee of Kyowa Kirin Co., Ltd., an AAALAC-accredited institution. Male C57BL/6J mice (7 weeks old; Jackson Laboratory Japan) were used in this study. Two inhibitory anti-ADAMTS13 antibodies (scFv-1-4-16-Fc and scFv-1-4-20-Fc)^32^ were administered to mice at 5 mg/kg, or mice received vehicle (Dulbecco’s phosphate-buffered saline [D-PBS]) at 5 mL/kg. After 24 h, ADAMTS13 mimetics were administered intravenously at 1-10 mg/kg, or acetate buffer vehicle (50 mM sodium acetate, 250 mM *D*-sorbitol, 50 mM *L*-arginine, 7.15 µg/mL ZnSO_4_, pH 5.0) was given at 5 mL/kg. Within 5 min of this dosing, the mice received an additional tail vein injection of recombinant VWF (rVWF; 500 or 600 U/kg) or its formulation buffer. After 24 h, 630 µL of blood was collected using a syringe preloaded with 70 µL of 3.2% sodium citrate buffer (pH 5.5). Blood was gently mixed, and 170 µL was transferred to EDTA-containing tubes for platelet counting. Platelet counts were measured using an automated hematology analyzer (XN-2000, Sysmex, Kobe, Japan). The remaining blood was centrifuged at 800 g for 10 min at 4°C, and plasma was collected for measurement of ADAMTS13 activity. For the fluorogenic ADAMTS13 assay, plasma samples were diluted to 15% (v/v) in FRET buffer, and recombinant ADAMTS13 was used as a standard. Plasma ADAMTS13 activity was calculated as a recombinant ADAMTS13-equivalent concentration based on FRETS-VWF73 cleavage and expressed as relative activity normalized to the saline-treated control group.

## Results

### Protein expression and purification of ADAMTS13 mimetics comprising the ADAMTS13 metalloprotease domain fused to anti-VWF A2 domain antibodies

The ADAMTS13 metalloproteinase domain alone has been reported to be insufficient to degrade VWF because of its limited binding activity to VWF.^6,33^ Therefore, we hypothesized that ADAMTS13 activity could be restored by fusing the ADAMTS13 metalloproteinase domain to an antibody that binds to the VWF A2 domain, thereby recruiting the catalytic domain to the VWF cleavage site (Figure 1A). We hereafter refer to these engineered fusion proteins as ADAMTS13 mimetics.

**Figure 1.**
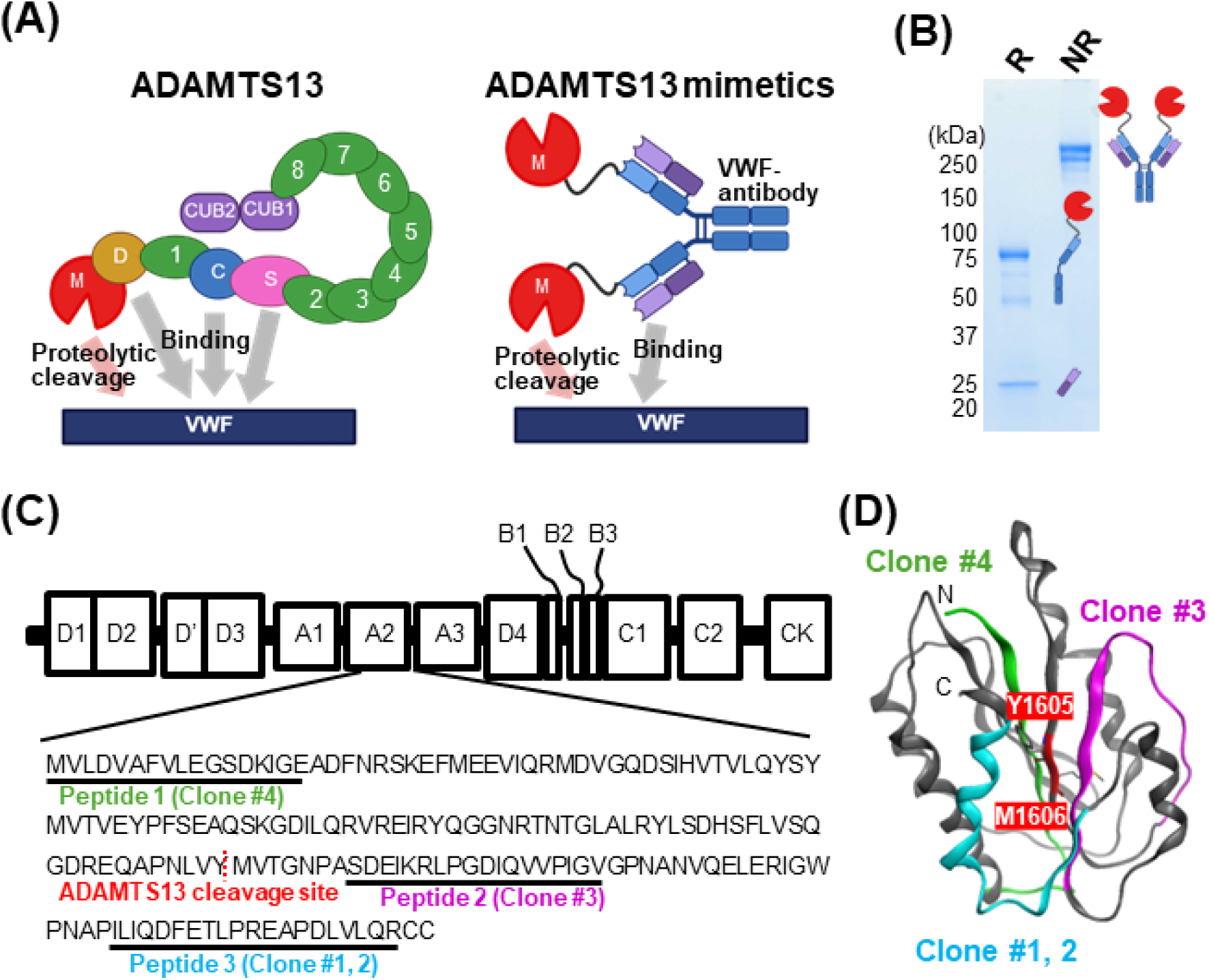
Design and purification of ADAMTS13 mimetics comprising the ADAMTS13 metalloproteinase domain fused to anti-VWF A2 domain antibodies. (A) Schematic representation of the ADAMTS13 mimetic design. The ADAMTS13 metalloproteinase domain was fused to an anti-VWF A2 domain antibody to recruit the catalytic domain to the VWF A2 domain and restore proteolytic activity toward VWF. (B) Representative SDS-PAGE analysis of purified ADAMTS13 mimetic #1. R, reducing conditions; NR, non-reducing conditions. (C) Schematic representation of the VWF domain organization and the positions of the immunizing peptides within the VWF A2 domain. The ADAMTS13 cleavage site, Tyr1605-Met1606, and the three immunizing peptide regions are indicated. Peptide 1, Peptide 2, and Peptide 3 are shown in green, magenta, and cyan, respectively. (D) Mapping of the designed peptides onto the crystal structure of the VWF A2 domain (PDB: 3GXB^13^). The immunizing peptide regions corresponding to the antibodies for Clone #4, Clone #3, and Clone #1-2 are highlighted in green, magenta, and cyan, respectively. The ADAMTS13 cleavage site is shown in red.

To test this concept, we generated antibodies using peptides designed around the ADAMTS13 cleavage site in the human VWF A2 domain, Tyr1605-Met1606. Three peptides corresponding to distinct regions near the cleavage site were used to immunize Kirin-Medarex (KM) humanized mice,^34^ and antibody clones binding to the respective immunizing peptide regions were obtained. The positions of the immunizing peptides were mapped onto both the VWF domain organization and the crystal structure of the VWF A2 domain (Figure 1C-D). Four ADAMTS13 mimetics, designated ADAMTS13 mimetics #1 to #4 according to their parental antibody clones, were generated from the four obtained anti-VWF A2 domain antibodies by fusing the ADAMTS13 metalloproteinase domain to the N terminus of each antibody heavy chain through a (GGGGS)_4_R linker (Figure 1A).

All four ADAMTS13 mimetics were transiently expressed in Expi293F cells and purified from culture supernatants by Protein A affinity chromatography. The purified proteins were analyzed by SDS-PAGE and size-exclusion chromatography to assess their purity (Figure 1B; Supplemental Figure 1). These analyses confirmed that ADAMTS13 metalloproteinase domain-anti-VWF A2 antibody fusion proteins were successfully produced as recombinant proteins suitable for subsequent biochemical characterization.

### *In vitro* binding and VWF A2-cleaving activity of ADAMTS13 mimetics

We next characterized the *in vitro* binding and VWF A2-cleaving activity of the ADAMTS13 mimetics. Binding to recombinant VWF A2 domain was evaluated by SPR. As a reference, ADAMTS13 bound to VWF A2 with high affinity (equilibrium dissociation constant [*K*_D_] = 24 nM; Supplemental Figure 2). All four ADAMTS13 mimetics also bound to VWF A2, with *K*_D_ values comparable to, or lower than, those of ADAMTS13 and the corresponding parental antibodies (Figure 2A; Table 1). These results indicate that fusion of the ADAMTS13 metalloproteinase domain to the antibody heavy chain preserved VWF A2 binding and enabled VWF A2 engagement with affinity comparable to that of the native ADAMTS13 substrate-recognition region.

**Figure 2.**
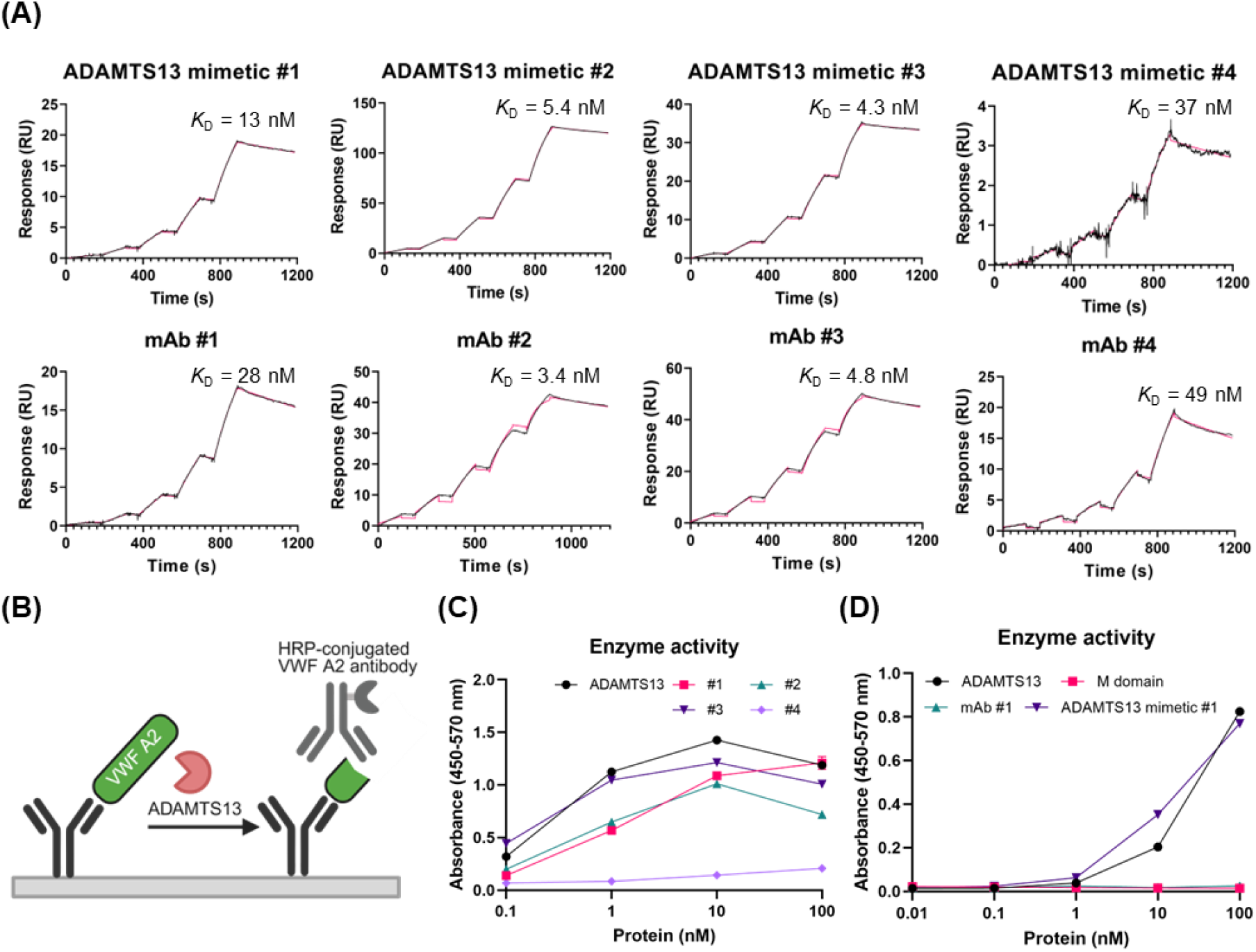
*In vitro* binding and VWF A2-cleaving activity of ADAMTS13 mimetics. (A) SPR analysis of ADAMTS13 mimetics and the corresponding parental anti-VWF A2 antibodies. Representative sensorgrams from three independent biological replicates are shown. Black lines indicate raw sensorgrams, and red lines indicate fitted curves. (B) Schematic illustration of the ELISA-based VWF A2 cleavage assay. (C) Concentration-dependent VWF A2-cleaving activity of ADAMTS13 and ADAMTS13 mimetics measured using the ELISA-based cleavage assay. Plates were coated with anti-His antibody at 5 µg/mL, and the N-terminal His_8_-tagged recombinant VWF A2 domain was captured at 2 µg/mL. The immobilized VWF A2 domain was incubated with ADAMTS13 or ADAMTS13 mimetics for 5 h. Absorbance was measured at 450 nm with background correction at 570 nm. Data are shown as mean ± SEM from three biological replicates. SEM, standard error of the mean. (D) Comparison of VWF A2-cleaving activity among ADAMTS13 mimetic #1, ADAMTS13 metalloproteinase domain alone, and parental anti-VWF A2 antibody. Plates were coated with anti-RFP antibody at 5 µg/mL, and the mCherry-tagged VWF A2 domain was immobilized at 2 µg/mL. The immobilized VWF A2 domain was incubated with ADAMTS13 mimetic, ADAMTS13 metalloproteinase domain alone, or parental anti-VWF A2 antibody for 3 h. Cleavage of the VWF A2 domain was detected using an HRP-labeled antibody recognizing the ADAMTS13-cleaved VWF A2 domain. Data are shown as mean ± SEM from three biological replicates.

**Table 1.** Binding kinetics of ADAMTS13 mimetics and parental anti-VWF A2 antibodies to recombinant VWF A2 domains.

| Protein | $k_a$, $M^{-1} s^{-1}$ | $k_d$, $s^{-1}$ | $K_D$, nM |
| --- | --- | --- | --- |
| ADAMTS13 mimetic #1 | $(1.90 \pm 0.27) \times 10^4$ | $(2.44 \pm 0.56) \times 10^{-4}$ | $12.9 \pm 2.0$ |
| Parental mAb #1 | $(1.78 \pm 0.30) \times 10^4$ | $(4.84 \pm 0.29) \times 10^{-4}$ | $27.7 \pm 3.5$ |
| ADAMTS13 mimetic #2 | $(3.11 \pm 0.56) \times 10^4$ | $(1.63 \pm 0.12) \times 10^{-4}$ | $5.43 \pm 1.45$ |
| Parental mAb #2 | $(7.34 \pm 0.59) \times 10^4$ | $(2.51 \pm 0.06) \times 10^{-4}$ | $3.43 \pm 0.27$ |
| ADAMTS13 mimetic #3 | $(3.69 \pm 0.37) \times 10^4$ | $(1.57 \pm 0.04) \times 10^{-4}$ | $4.28 \pm 0.49$ |
| Parental mAb #3 | $(6.34 \pm 0.65) \times 10^4$ | $(3.01 \pm 0.07) \times 10^{-4}$ | $4.80 \pm 0.56$ |
| ADAMTS13 mimetic #4 | $(1.53 \pm 1.01) \times 10^4$ | $(4.80 \pm 2.57) \times 10^{-4}$ | $36.5 \pm 15.2$ |
| Parental mAb #4 | $(1.33 \pm 0.22) \times 10^4$ | $(6.42 \pm 0.33) \times 10^{-4}$ | $48.9 \pm 5.2$ |

Values are shown as mean ± SD from three independent measurements for A2-WT binding.
$k_a$ indicates the association rate constant; $k_d$ , the dissociation rate constant; $K_D$ , the equilibrium dissociation constant ( $K_D = k_d/k_a$ ); mAb, monoclonal antibody; and SD, standard deviation.

We then examined whether the ADAMTS13 mimetics could cleave the VWF A2 domain using an ELISA-based cleavage assay (Figure 2B). In this assay, a plate-immobilized recombinant VWF A2 domain was incubated with ADAMTS13 or ADAMTS13 mimetics, and VWF A2 cleavage was detected using an antibody that recognizes the ADAMTS13-cleaved VWF A2 domain. ADAMTS13 mimetics #1, #2, and #3 showed concentration-dependent VWF A2-cleaving activity, whereas ADAMTS13 mimetic #4 showed markedly lower activity (Figure 2C).

To determine whether VWF A2 cleavage required both the metalloproteinase domain and antibody-mediated VWF A2 recognition, we compared the activity of an ADAMTS13 mimetic with that of its individual components. Neither the ADAMTS13 metalloproteinase domain alone nor the parental anti-VWF A2 antibody showed detectable VWF A2-cleaving activity, whereas the corresponding ADAMTS13 mimetic showed concentration-dependent cleavage activity (Figure 2D). These results indicate that fusion of the ADAMTS13 metalloproteinase domain to an anti-VWF A2 antibody reconstitutes the VWF A2-cleaving activity, and that both the catalytic domain and VWF A2-targeting antibody moiety are required for efficient cleavage.

### Enzyme kinetic characterization of ADAMTS13 mimetics using FRETS-VWF73

To further evaluate the catalytic properties of ADAMTS13 mimetics, enzyme kinetic analysis was performed using the fluorogenic FRETS-VWF73 substrate.^35^ We first compared full-length ADAMTS13 with ADAMTS13 mimetic #1 by monitoring fluorescence signals every 5 min for 60 min over a range of FRETS-VWF73 concentrations. Both ADAMTS13 and ADAMTS13 mimetic #1 showed time-dependent FRETS-VWF73 cleavage (Figure 3A). Initial reaction velocities were calculated from these fluorescence time-course data and used to determine kinetic parameters. ADAMTS13 mimetic #1 showed a lower Michaelis constant (*K*_m_) than ADAMTS13 (10.7 µM and 22.8 µM, respectively; Figure 3B). In contrast, the turnover number (*k*_cat_) of ADAMTS13 mimetic #1 was lower than that of ADAMTS13 (2.79 min^−1^ and 13.5 min^−1^, respectively; Figure 3C). Despite the reduction in *k*_cat_, ADAMTS13 mimetic #1 maintained approximately 44% of the apparent catalytic efficiency of ADAMTS13, as assessed by *k*_cat_/*K*_m_ (Figure 3D). These results indicate that ADAMTS13 mimetic #1 retained substantial FRETS-VWF73-cleaving activity because its reduced catalytic turnover was offset by a lower *K*_m_.

**Figure 3.**
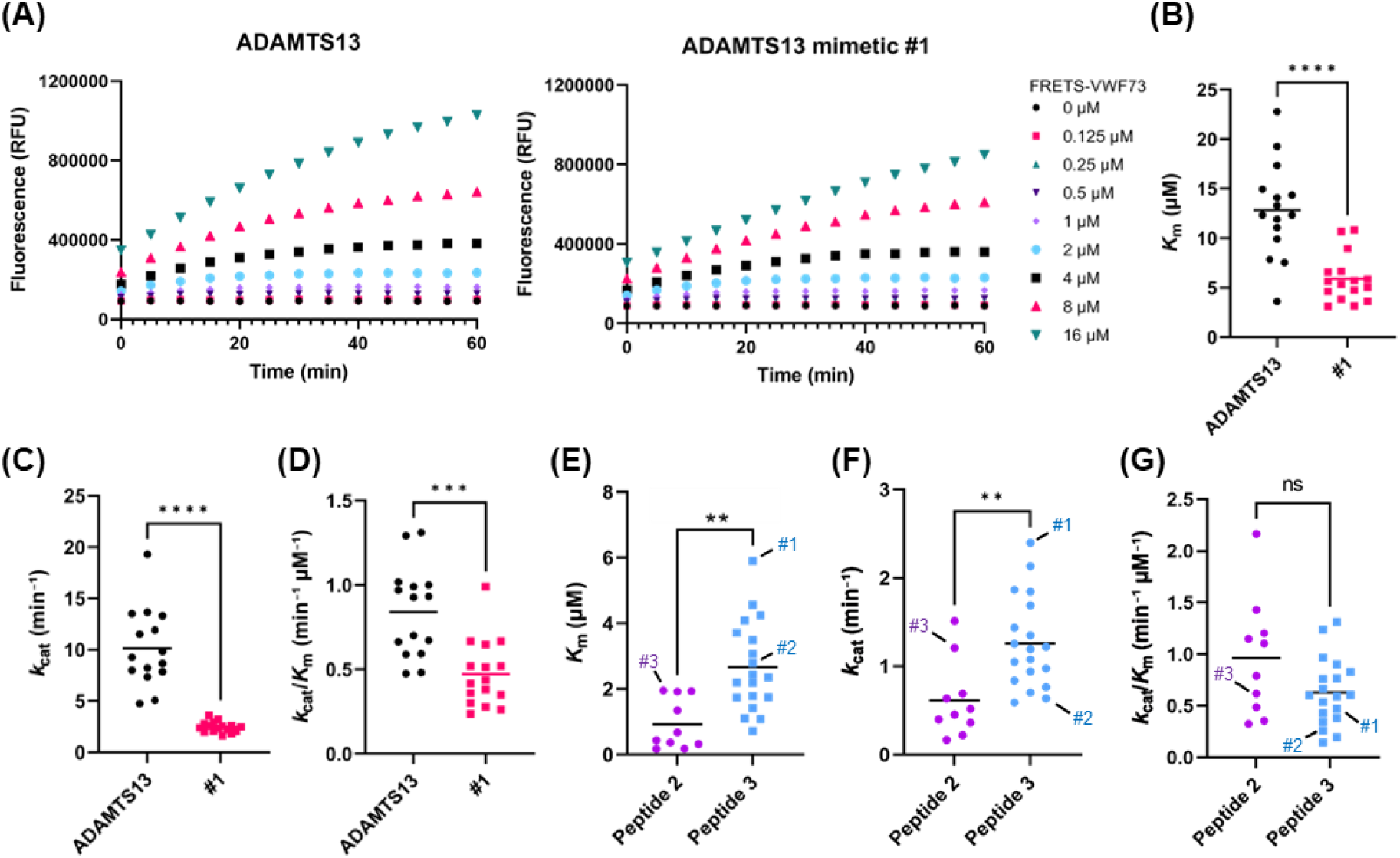
Enzyme kinetic characterization of ADAMTS13 mimetics using FRETS-VWF73. (A) Representative fluorescence time-course data for FRETS-VWF73 cleavage by ADAMTS13 and ADAMTS13 mimetic #1. Fluorescence signals were measured every 5 min for 60 min over a range of FRETS-VWF73 concentrations. ADAMTS13 and ADAMTS13 mimetic #1 were used at final concentrations of 30 nM and 100 nM, respectively. (B-D) Enzyme kinetic analysis of ADAMTS13 and ADAMTS13 mimetic #1 using FRETS-VWF73. Each point represents the kinetic parameter calculated from one independent biological replicate (ADAMTS13, *n* = 15; ADAMTS13 mimetic #1, *n* = 16). The *K*_m_ (B), *k*_cat_ (C), and *k*_cat_/*K*_m_ (D) are shown. Horizontal bars indicate mean values. Statistical significance was determined using a two-tailed Mann-Whitney test with Holm-Bonferroni correction. \*\*\**P* < 0.001; \*\*\*\**P* < 0.0001. (E-G) Kinetic comparison of additional ADAMTS13 mimetics derived from antibodies recognizing Peptide 2 or Peptide 3. Each point represents one ADAMTS13 mimetic clone (Peptide 2, *n* = 10; Peptide 3, *n* = 19). Apparent *K*_m_ (E), *k*_cat_ (F), and *k*_cat_/*K*_m_ (G) are shown. Horizontal bars indicate mean values. Statistical significance was determined using a two-tailed Mann-Whitney test with Holm-Bonferroni correction. \*\**P* < 0.01; ns, not significant.

Given that ADAMTS13 mimetics targeting Peptide 2 and Peptide 3 showed higher activity in the VWF A2 cleavage assay compared with Peptide 1 (Figures 1C and 2B-C), we further evaluated additional metalloproteinase domain fusion constructs derived from antibodies recognizing these peptides. Peptide 3-derived mimetics showed significantly lower apparent *K*_m_ values than Peptide 2-derived mimetics (Figure 3E). Peptide 3-derived mimetics also showed significantly higher apparent *k*_cat_ values than Peptide 2-derived mimetics (Figure 3F). However, apparent *k*_cat_/*K*_m_ was not significantly different between the two groups (Figure 3G). Notably, several ADAMTS13 mimetics exhibited apparent *k*_cat_/*K*_m_ values in the range of full-length ADAMTS13, although this comparison was limited by small sample sizes (Supplemental Figure 3).

Overall, these results indicate that the antibody-binding region targeted within the VWF A2 domain influences both substrate engagement and catalytic turnover of ADAMTS13 mimetics. Although the engineered fusion format reduced the catalytic turnover in ADAMTS13 mimetic #1, the lower *K*_m_ preserved catalytic efficiency. In addition, several ADAMTS13 mimetics exhibited apparent catalytic efficiencies within the range of full-length ADAMTS13. These findings suggest that antibody-guided positioning of the ADAMTS13 metalloproteinase domain can preserve overall enzymatic function toward FRETS-VWF73.

### ADAMTS13 mimetics prevent thrombocytopenia in iTTP and ADAMTS13-deficient mouse models

We next evaluated whether ADAMTS13 mimetics could function under conditions that mimic iTTP. We first examined whether ADAMTS13 mimetics retained their VWF A2-cleaving activity in the presence of the spacer-directed inhibitory anti-ADAMTS13 scFv 1-4-20.^32,36^ Full-length ADAMTS13 and recombinant ADAMTS13_MDTCS, which contains the metalloproteinase, disintegrin-like, thrombospondin type 1, cysteine-rich, and spacer domains, were strongly inhibited by scFv 1-4-20 (Figure 4A). In contrast, ADAMTS13 mimetics retained the VWF A2-cleaving activity under the same conditions (Figure 4A). These results indicate that ADAMTS13 mimetics can bypass inhibition by a spacer-directed anti-ADAMTS13 antibody.

**Figure 4.**
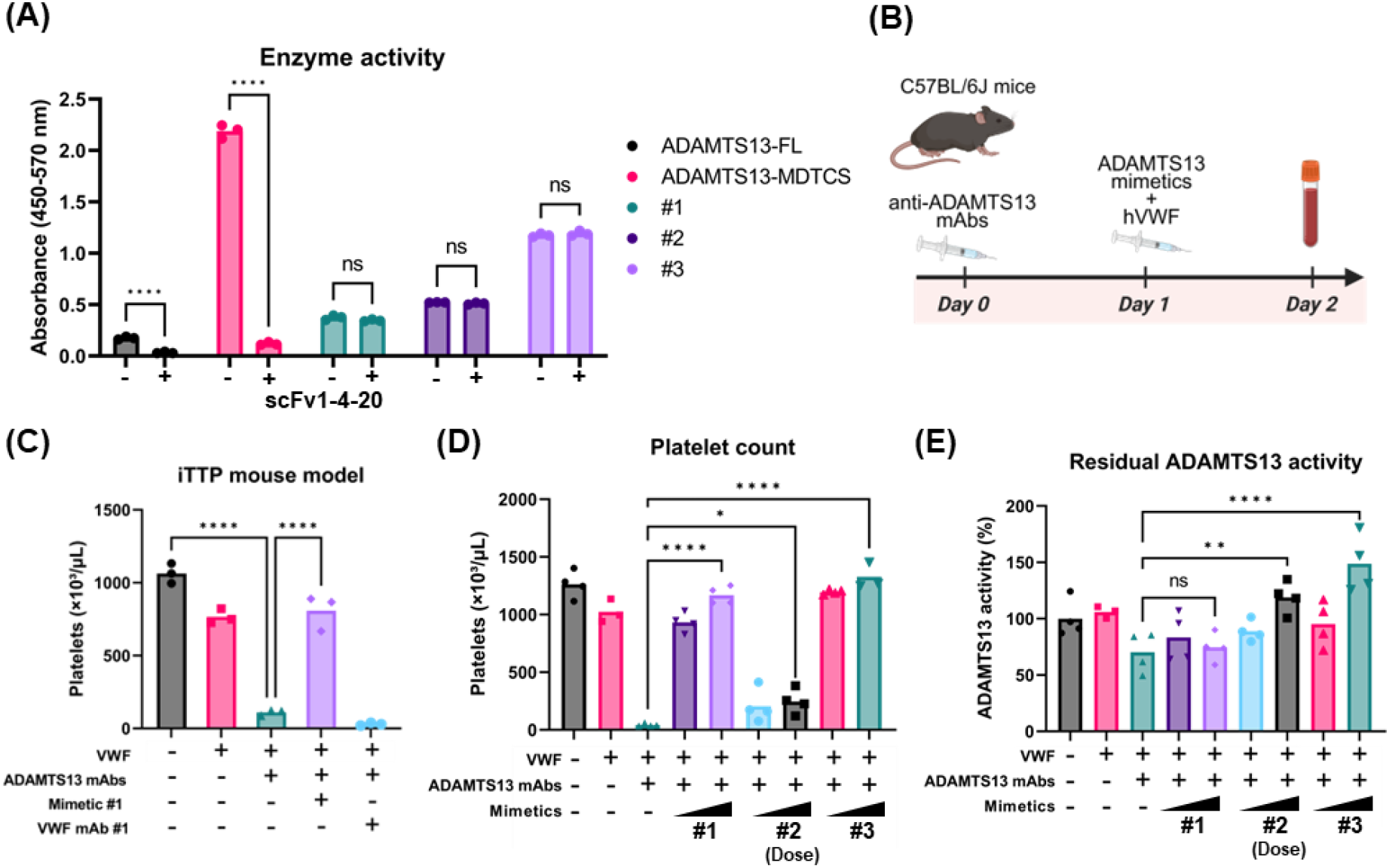
ADAMTS13 mimetics prevent thrombocytopenia in an iTTP mouse model. (A) Effect of the patient-derived anti-ADAMTS13 scFv 1-4-20 on VWF A2-cleaving activity. The mCherry-tagged VWF A2 domain was captured on anti-RFP antibody-coated plates and incubated for 3 h with full-length ADAMTS13, ADAMTS13_MDTCS, or ADAMTS13 mimetics in the presence or absence of scFv 1-4-20. ADAMTS13 proteins and scFv 1-4-20 were used at final concentrations of 5 nM and 25 nM, respectively. VWF A2-cleaving activity was measured by ELISA using an HRP-labeled antibody recognizing the ADAMTS13-cleaved VWF A2 domain. Data are shown as mean from three biological replicates. Statistical significance was determined using a two-way ANOVA, followed by Šídák’s multiple-comparisons test comparing conditions with and without scFv 1-4-20 within each protein. \*\*\*\**P* < 0.0001; ns, not significant. (B) Schematic representation of the iTTP mouse model. Mice were treated intravenously with inhibitory anti-ADAMTS13 antibodies on day 0. On day 1, mice received recombinant human VWF together with a vehicle, parental anti-VWF A2 antibody, or ADAMTS13 mimetic. Platelet counts and residual plasma ADAMTS13 activity were measured on day 2. (C) Platelet counts in mice treated with vehicle, recombinant human VWF, inhibitory anti-ADAMTS13 antibodies, parental anti-VWF A2 antibody, or ADAMTS13 mimetic #1. The inhibitory anti-ADAMTS13 antibody mixture, consisting of scFv 1-4-20-Fc and scFv 1-4-16-Fc, was administered at a total dose of 5 mg/kg. Recombinant human VWF was administered at 600 U/kg, ADAMTS13 mimetic #1 at 10 mg/kg, and the parental anti-VWF A2 antibody at 10 mg/kg. Data are shown as mean, *n* = 3 mice per group. Statistical significance was determined by a one-way ANOVA with Dunnett’s multiple-comparisons test. \*\*\*\**P* < 0.0001. (D) Comparison of platelet-preserving activity among ADAMTS13 mimetics #1, #2, and #3. The inhibitory anti-ADAMTS13 antibody mixture, consisting of scFv 1-4-20-Fc and scFv 1-4-16-Fc, was administered at a total dose of 5 mg/kg. Recombinant human VWF was administered at 500 U/kg, and ADAMTS13 mimetics were administered at 1 or 5 mg/kg. Data are shown as mean, *n* = 4 mice per group. Statistical significance was determined by a one-way ANOVA with Dunnett’s multiple-comparisons test unless otherwise indicated. \**P* < 0.05; \*\*\*\**P* < 0.0001. (E) Residual plasma ADAMTS13 activity measured using the FRETS-VWF73 substrate. Residual activity was normalized to the activity detected in the saline-treated control group. *n* = 4 mice per group. Data are shown as mean. Statistical significance was determined by a one-way ANOVA with Dunnett’s multiple-comparisons test unless otherwise indicated. \*\**P* < 0.01; \*\*\*\**P* < 0.0001; ns, not significant.

We then assessed the therapeutic activity of ADAMTS13 mimetics in an iTTP mouse model. Mice were administered inhibitory anti-ADAMTS13 antibodies, followed by recombinant human VWF together with a vehicle, the parental anti-VWF A2 antibody, or an ADAMTS13 mimetic. Platelet counts and residual plasma ADAMTS13 activity were evaluated 24 h after the VWF challenge (Figure 4B). Recombinant VWF administration in anti-ADAMTS13 antibody-treated mice induced marked thrombocytopenia (Figure 4C). Treatment with the parental anti-VWF A2 antibody did not restore platelet counts, indicating that VWF A2 binding alone was insufficient for protection. In contrast, ADAMTS13 mimetic treatment restored platelet counts in this model (Figure 4C). These results demonstrate that the metalloproteinase domain fused to a VWF A2-targeting antibody is required to prevent thrombocytopenia *in vivo*.

To assess whether this platelet-preserving activity was independent of endogenous ADAMTS13, we tested the mimetics in ADAMTS13-deficient mice.^37^ Intravenous administration of recombinant human VWF induced marked thrombocytopenia in vehicle-treated ADAMTS13-deficient mice (Supplemental Figure 4A). Treatment with either ADAMTS13 mimetic #1 or #3 restored platelet counts, as did recombinant ADAMTS13 (Supplemental Figure 4A-B). These results confirm that ADAMTS13 mimetics can prevent VWF-induced thrombocytopenia *in vivo* even in the absence of endogenous ADAMTS13.

We then compared the *in vivo* efficacy of multiple ADAMTS13 mimetic clones in an iTTP mouse model. ADAMTS13 mimetics #1, #2, and #3 increased platelet counts compared with vehicle-treated disease-control mice, although the magnitude of platelet recovery differed among clones (Figure 4D). To determine whether platelet recovery was associated with circulating enzymatic activity, we measured residual plasma ADAMTS13 activity using the FRETS-VWF73 substrate. Residual activity differed among mimetics and did not fully parallel the platelet-preserving effects observed *in vivo* (Figure 4E). These findings suggest that the *in vivo* efficacy of ADAMTS13 mimetics is not determined solely by residual plasma activity measured with a soluble peptide substrate but may also depend on additional factors such as VWF epitope accessibility and productive positioning of the metalloproteinase domain on VWF.

Together, these results show that ADAMTS13 mimetics retain the VWF-cleaving activity in the presence of an inhibitory anti-ADAMTS13 antibody and prevent VWF-induced thrombocytopenia in both iTTP and ADAMTS13-deficient mouse models.

### Binding to conformationally restricted VWF A2 is associated with platelet-preserving activity of ADAMTS13 mimetics

The discordance between residual plasma ADAMTS13 activity and platelet protection suggested that the *in vivo* activity of ADAMTS13 mimetics is not explained solely by their catalytic activity toward the FRETS-VWF73 substrate. Therefore, we examined whether the conformational state of the VWF A2 domain and the antibody epitope recognized by each mimetic are associated with proteolytic activity and *in vivo* efficacy. Under physiological conditions, the VWF A2 domain adopts a folded conformation that limits access to the Tyr1605-Met1606 scissile bond. Mechanical unfolding exposes this cleavage site and enables proteolysis by ADAMTS13. To model a conformationally restricted A2 domain, we used a disulfide-locked VWF A2 variant (A2-locked) corresponding to the N1493C/C1670S mutation^38,39^ (Figure 5A). Consistent with the reported crystal structure showing preservation of the overall wild-type A2 fold (A2-WT), A2-locked was used to restrict conformational opening of A2 without globally disrupting the domain structure.^39^ However, in the ELISA-based VWF A2 cleavage assay, cleavage of A2-locked was markedly reduced compared with A2-WT for both ADAMTS13 and ADAMTS13 mimetics (Supplemental Figure 5). These results indicate that exposure of the scissile bond remains required for proteolysis by ADAMTS13 mimetics.

**Figure 5.**
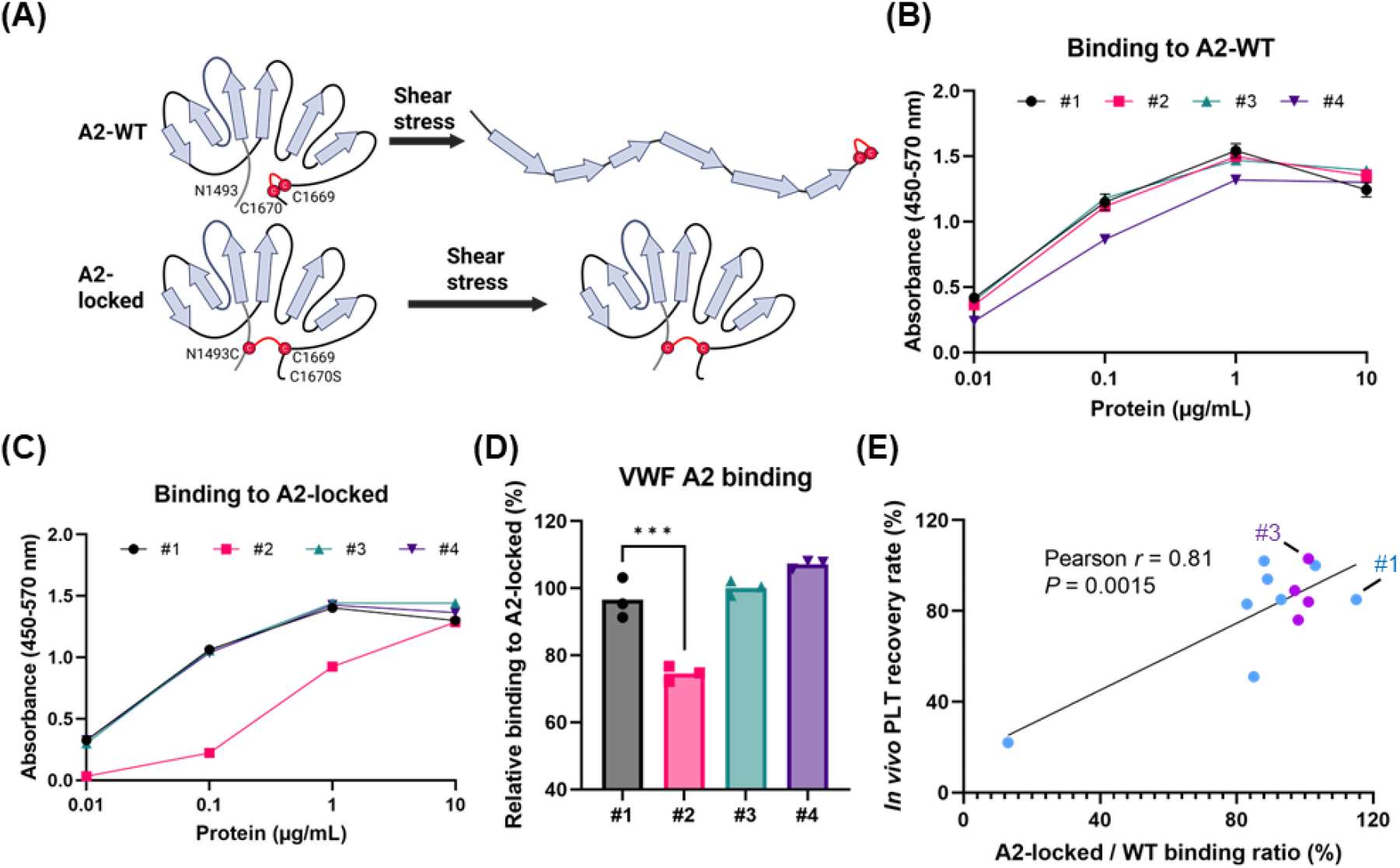
Relative binding to conformationally restricted VWF A2 is associated with platelet-preserving activity of ADAMTS13 mimetics. (A) Schematic representation of VWF A2 conformational regulation and the disulfide-locked A2 variant. Mechanical unfolding of A2 exposes the Tyr1605-Met1606 cleavage site, whereas A2-locked restricts conformational opening through an engineered disulfide bond. (B, C) Recombinant A2-WT or A2-locked was immobilized on plates at 2 µg/mL and incubated with serial dilutions of ADAMTS13 mimetics. Bound ADAMTS13 mimetics were detected using an HRP-conjugated anti-human IgG Fc antibody. Binding to A2-WT (B) and A2-locked (C) is shown. Data are shown as mean ± SEM from three biological replicates. (D) Relative binding of ADAMTS13 mimetics to conformationally restricted A2. The A2-locked/A2-WT binding ratio was calculated from the AUC values of the binding curves shown in panels B and C. Data are shown as mean from three biological replicates. Statistical significance was determined by a one-way ANOVA with Dunnett’s multiple-comparisons test. \*\*\**P* < 0.001 (E) Relationship between the A2-locked/A2-WT binding ratio and platelet recovery in the iTTP mouse model after the recombinant human VWF challenge. The A2-locked/A2-WT binding ratio was determined using parental monoclonal antibodies by ELISA-based binding assays from two independent experiments. Platelet recovery was evaluated using the corresponding ADAMTS13 mimetics in mice treated with the inhibitory anti-ADAMTS13 antibody mixture at a total dose of 30 mg/kg, recombinant human VWF at 500 U/kg, and ADAMTS13 mimetics at 5 mg/kg. Platelet recovery is shown as the mean value from four mice per group. Each point represents one antibody clone. Pearson correlation analysis was performed, and the line indicates the linear regression fit.

To determine whether differential recognition of conformationally restricted A2 could contribute to clone-dependent *in vivo* activity, we evaluated binding of ADAMTS13 mimetics to A2-WT and A2-locked. ELISA-based binding assays showed that ADAMTS13 mimetics bound to A2-WT in a concentration-dependent manner (Figure 5B). In contrast, binding to A2-locked differed among mimetics (Figure 5C). The A2-locked/A2-WT binding ratio was then calculated from the AUC values of the binding curves to quantify the relative recognition of conformationally restricted A2 (Figure 5D). Although mimetics #1 and #2 recognized the same epitope group, mimetic #1 showed a higher A2-locked/A2-WT binding ratio than mimetic #2 (Figure 5D). This difference was consistent with their *in vivo* activities, as mimetic #1 showed stronger platelet-preserving activity than mimetic #2 (Figure 4D). These results suggest that relative binding to conformationally restricted A2 may help explain the differences in *in vivo* activity among mimetics recognizing similar epitope regions.

To further examine this relationship, we compared the A2-locked/A2-WT binding ratio with the platelet recovery rate *in vivo*. Across Peptide 2- and Peptide 3-derived clones, the A2-locked/A2- WT binding ratio was positively associated with platelet recovery induced by the corresponding ADAMTS13 mimetics in the iTTP mouse model after the recombinant human VWF challenge (Figure 5E; Supplemental Figure 6). These findings suggest that recognition of epitopes accessible in conformationally restricted VWF A2 is associated with platelet-preserving activity *in vivo*.

Overall, these results indicate that ADAMTS13 mimetic activity is not determined by epitope assignment alone. Relative recognition of conformationally restricted A2 was associated with platelet-preserving activity *in vivo*, whereas reduced cleavage of A2-locked indicates that efficient proteolysis still requires exposure of the Tyr1605-Met1606 scissile bond. Thus, both accessibility of antibody-binding regions in a conformationally restricted VWF A2 domain and productive positioning of the metalloproteinase domain may contribute to optimal ADAMTS13 mimetic activity.

## Discussion

In this study, we developed ADAMTS13 mimetics by fusing the ADAMTS13 metalloproteinase domain to VWF A2-targeting antibodies. These constructs were designed to replace the substrate-recognition function of the non-catalytic ADAMTS13 domains with antibody-mediated targeting. The resulting mimetics bound to VWF A2, cleaved VWF-derived substrates, bypassed inhibition by a spacer-directed anti-ADAMTS13 autoantibody, and prevented VWF-induced thrombocytopenia in iTTP and ADAMTS13-deficient mouse models. Given that inhibitory autoantibodies in iTTP frequently target the spacer and adjacent substrate-recognition domains, removal of these domains may reduce susceptibility to autoantibody inhibition.^26,27,40^ Thus, ADAMTS13 mimetics may provide a potential strategy for autoantibody-resistant ADAMTS13 replacement therapy.

A key finding is that VWF A2 binding alone is not sufficient for efficient cleavage. The Peptide 1-derived mimetic showed low activity, whereas multiple Peptide 2- or Peptide 3-derived mimetics showed robust VWF-cleaving activity. These results suggest that productive mimetic activity requires antibody binding to regions that enable effective recruitment and spatial alignment of the metalloproteinase domain relative to the Tyr1605-Met1606 cleavage site. Kinetic analysis supported this model. Most mimetics showed lower *k*_cat_ values than ADAMTS13, but several maintained overall catalytic efficiency through lower *K*_m_ values. Thus, ADAMTS13 mimetics appear to restore VWF cleavage mainly through antibody-mediated substrate targeting rather than by fully reproducing native exosite-dependent activation.

The conformational state of VWF A2 also influenced mimetic activity. A2-locked was used as a previously characterized conformationally restricted A2 variant. Cleavage of A2-locked was markedly reduced, indicating that exposure of the Tyr1605-Met1606 scissile bond remains required for proteolysis. At the same time, relative recognition of A2-locked was associated with platelet-preserving activity *in vivo*. These findings suggest that both epitope accessibility in conformationally restricted VWF A2 and productive positioning of the metalloproteinase domain contribute to optimal ADAMTS13 mimetic activity.

VWF-directed enzymatic therapy has also been explored using Microlyse, a VWF-targeting variable domain of a heavy-chain-only antibody (VHH) fused to the urokinase-type plasminogen activator (uPA) protease domain, which attenuated thrombocytopenia in an ADAMTS13-deficient mouse model.^41^ Microlyse promotes plasmin-mediated degradation of VWF-rich microthrombi, whereas our ADAMTS13 mimetics were designed to reconstitute ADAMTS13-like cleavage of the VWF A2 domain. Thus, our findings extend this concept by showing that antibody-mediated substrate targeting can restore VWF-cleaving activity through an ADAMTS13-based mechanism. In conclusion, antibody-mediated targeting of the ADAMTS13 metalloproteinase domain to VWF A2 reconstituted the VWF-cleaving activity and prevented VWF-induced thrombocytopenia *in vivo*. These findings establish ADAMTS13 mimetics as a potential therapeutic approach for iTTP and provide a framework for further optimization of antibody-guided VWF-cleaving enzymes.

## Acknowledgments

The authors thank the Laboratory of Animal Models for Human Diseases, National Institutes of Biomedical Innovation, Health and Nutrition, Japan, for providing frozen embryos of ADAMTS13 knockout mice. The authors thank members of the Research Division at Kyowa Kirin for helpful discussions and technical advice throughout the study. Editorial assistance, limited to English-language editing, was provided by Cactus Life Sciences and was funded by Kyowa Kirin Co., Ltd. The visual abstract and schematic illustrations were created with BioRender.com under a publication license.

## Authorship

### Contribution

**A.H**.: Conceptualization, Investigation, Visualization, Writing - original draft. **Y.K**.: Investigation, Visualization. **F.K**.: Investigation. **M.O**.: Investigation. **M.N**.: Investigation. **S.S**.: Investigation. **H.I**.: Supervision. **Y.S**.: Supervision. **K.M**.: Conceptualization, Project administration, Supervision. **A.S**.: Conceptualization, Writing - review & editing, Project administration, Supervision.

## References

1. Zheng X, Chung D, Takayama TK, Majerus EM, Sadler JE, Fujikawa K. Structure of von Willebrand factor-cleaving protease (ADAMTS13), a metalloprotease involved in thrombotic thrombocytopenic purpura. J Biol Chem. 2001;276(44):41059–41063.

2. Crawley JTB, de Groot R, Xiang Y, Luken BM, Lane DA. Unraveling the scissile bond: how ADAMTS13 recognizes and cleaves von Willebrand factor. Blood. 2011;118(12):3212–3221.

3. Zheng XL. Structure-function and regulation of ADAMTS-13 protease. J Thromb Haemost. 2013;11 Suppl 1(0 1):11–23.

4. Fujikawa K, Suzuki H, McMullen B, Chung D. Purification of human von Willebrand factor-cleaving protease and its identification as a new member of the metalloproteinase family. Blood. 2001;98(6):1662–1666.

5. Zander CB, Cao W, Zheng XL. ADAMTS13 and von Willebrand factor interactions. Curr Opin Hematol. 2015;22(5):452–459.

6. Ai J, Smith P, Wang S, Zhang P, Zheng XL. The proximal carboxyl-terminal domains of ADAMTS13 determine substrate specificity and are all required for cleavage of von Willebrand factor. J Biol Chem. 2005;280(33):29428–29434.

7. Gao W, Anderson PJ, Majerus EM, Tuley EA, Sadler JE. Exosite interactions contribute to tension-induced cleavage of von Willebrand factor by the antithrombotic ADAMTS13 metalloprotease. Proc Natl Acad Sci U S A. 2006;103(50):19099–19104.

8. Gao W, Zhu J, Westfield LA, Tuley EA, Anderson PJ, Sadler JE. Rearranging exosites in noncatalytic domains can redirect the substrate specificity of ADAMTS proteases. J Biol Chem. 2012;287(32):26944–26952.

9. Geist N, Nagel F, Delcea M. Molecular interplay of ADAMTS13-MDTCS and von Willebrand factor-A2: deepened insights from extensive atomistic simulations. J Biomol Struct Dyn. 2023;41(17):8201–8214.

10. Wu J-J, Fujikawa K, McMullen BA, Chung DW. Characterization of a core binding site for ADAMTS-13 in the A2 domain of von Willebrand factor. Proc Natl Acad Sci U S A. 2006;103(49):18470–18474.

11. Akiyama M, Takeda S, Kokame K, Takagi J, Miyata T. Crystal structures of the noncatalytic domains of ADAMTS13 reveal multiple discontinuous exosites for von Willebrand factor. Proc Natl Acad Sci U S A. 2009;106(46):19274–19279.

12. Petri A, Kim HJ, Xu Y, et al. Crystal structure and substrate-induced activation of ADAMTS13. Nat Commun. 2019;10(1):3781.

13. Zhang Q, Zhou Y-F, Zhang C-Z, Zhang X, Lu C, Springer TA. Structural specializations of A2, a force-sensing domain in the ultralarge vascular protein von Willebrand factor. Proc Natl Acad Sci U S A. 2009;106(23):9226–9231.

14. Lynch CJ, Lane DA, Luken BM. Control of VWF A2 domain stability and ADAMTS13 access to the scissile bond of full-length VWF. Blood. 2014;123(16):2585–2592.

15. Scully M, Hunt BJ, Benjamin S, et al. Guidelines on the diagnosis and management of thrombotic thrombocytopenic purpura and other thrombotic microangiopathies. Br J Haematol. 2012;158(3):323–335.

16. Joly BS, Coppo P, Veyradier A. Thrombotic thrombocytopenic purpura. Blood. 2017;129(21):2836–2846.

17. Kremer Hovinga JA, Coppo P, Lämmle B, Moake JL, Miyata T, Vanhoorelbeke K. Thrombotic thrombocytopenic purpura. Nat Rev Dis Primers. 2017;3:17020.

18. Joly BS, Stepanian A, Leblanc T, et al. Child-onset and adolescent-onset acquired thrombotic thrombocytopenic purpura with severe ADAMTS13 deficiency: a cohort study of the French national registry for thrombotic microangiopathy. Lancet Haematol. 2016;3(11):e537–e546.

19. Mariotte E, Azoulay E, Galicier L, et al. Epidemiology and pathophysiology of adulthood-onset thrombotic microangiopathy with severe ADAMTS13 deficiency (thrombotic thrombocytopenic purpura): a cross-sectional analysis of the French national registry for thrombotic microangiopathy. Lancet Haematol. 2016;3(5):e237–e245.

20. Fujimura Y, Matsumoto M. Registry of 919 patients with thrombotic microangiopathies across Japan: database of Nara Medical University during 1998-2008. Intern Med. 2010;49(1):7–15.

21. Scully M, Antun A, Cataland SR, et al. Recombinant ADAMTS13 in congenital thrombotic thrombocytopenic purpura. N Engl J Med. 2024;390(17):1584–1596.

22. Zheng XL. The standard of care for immune thrombotic thrombocytopenic purpura today. J Thromb Haemost. 2021;19(8):1864–1871.

23. Capecchi M, Gazzola G, Agosti P, et al. Treatment of immune-mediated thrombotic thrombocytopenic purpura without plasma exchange. Haematologica. 2024;109(6):2019–2023.

24. Klaus C, Plaimauer B, Studt J-D, et al. Epitope mapping of ADAMTS13 autoantibodies in acquired thrombotic thrombocytopenic purpura. Blood. 2004;103(12):4514–4519.

25. Velásquez Pereira LC, Roose E, Graça NAG, et al. Immunogenic hotspots in the spacer domain of ADAMTS13 in immune-mediated thrombotic thrombocytopenic purpura. J Thromb Haemost. 2021;19(2):478–488.

26. Tersteeg C, Verhenne S, Roose E, et al. ADAMTS13 and anti-ADAMTS13 autoantibodies in thrombotic thrombocytopenic purpura - current perspectives and new treatment strategies. Expert Rev Hematol. 2016;9(2):209–221.

27. Thomas MR, de Groot R, Scully MA, Crawley JTB. Pathogenicity of anti-ADAMTS13 autoantibodies in acquired thrombotic thrombocytopenic purpura. eBioMedicine. 2015;2(8):942–952.

28. Jian C, Xiao J, Gong L, et al. Gain-of-function ADAMTS13 variants that are resistant to autoantibodies against ADAMTS13 in patients with acquired thrombotic thrombocytopenic purpura. Blood. 2012;119(16):3836–3843.

29. Kwak H, Choi G, Kim S, et al. GC1126A, a novel ADAMTS13 mutein, evades autoantibodies in immune-mediated thrombotic thrombocytopenic purpura. Sci Rep. 2025;15(1):1613.

30. Ercig B, Graça NAG, Kangro K, et al. N-Glycan-mediated shielding of ADAMTS13 prevents binding of pathogenic autoantibodies in immune-mediated TTP. Blood. 2021;137(19):2694–2698.

31. Graça NAG, Ercig B, Carolina Velásquez Pereira L, et al. Modifying ADAMTS13 to modulate binding of pathogenic autoantibodies of patients with acquired thrombotic thrombocytopenic purpura. Haematologica. 2020;105(11):2619–2630.

32. Ostertag EM, Kacir S, Thiboutot M, et al. ADAMTS13 autoantibodies cloned from patients with acquired thrombotic thrombocytopenic purpura: 1. Structural and functional characterization in vitro. Transfusion. 2016;56(7):1763–1774.

33. Zheng X, Nishio K, Majerus EM, Sadler JE. Cleavage of von Willebrand factor requires the spacer domain of the metalloprotease ADAMTS13. J Biol Chem. 2003;278(32):30136–30141.

34. Ishida I, Tomizuka K, Yoshida H, et al. Production of human monoclonal and polyclonal antibodies in TransChromo animals. Cloning Stem Cells. 2002;4(1):91–102.

35. Kokame K, Nobe Y, Kokubo Y, Okayama A, Miyata T. FRETS-VWF73, a first fluorogenic substrate for ADAMTS13 assay. Br J Haematol. 2005;129(1):93–100.

36. Halkidis K, Meng C, Liu S, Mayne L, Siegel DL, Zheng XL. Mechanisms of inhibition of human monoclonal antibodies in immune thrombotic thrombocytopenic purpura. Blood. 2023;141(24):2993–3005.

37. Banno F, Kokame K, Okuda T, et al. Complete deficiency in ADAMTS13 is prothrombotic, but it alone is not sufficient to cause thrombotic thrombocytopenic purpura. Blood. 2006;107(8):3161–3166.

38. Baldauf C, Schneppenheim R, Stacklies W, et al. Shear-induced unfolding activates von Willebrand factor A2 domain for proteolysis. J Thromb Haemost. 2009;7(12):2096–2105.

39. Zhou M, Dong X, Baldauf C, et al. A novel calcium-binding site of von Willebrand factor A2 domain regulates its cleavage by ADAMTS13. Blood. 2011;117(17):4623–4631.

40. Zheng XL, Wu HM, Shang D, et al. Multiple domains of ADAMTS13 are targeted by autoantibodies against ADAMTS13 in patients with acquired idiopathic thrombotic thrombocytopenic purpura. Haematologica. 2010;95(9):1555–1562.

41. de Maat S, Clark CC, Barendrecht AD, et al. Microlyse: a thrombolytic agent that targets VWF for clearance of microvascular thrombosis. Blood. 2022;139(4):597–607.

