## Supplemental data for "Antibody-mediated VWF targeting reconstitutes ADAMTS13 metalloproteinase activity and prevents thrombocytopenia in mice"

**Procedure for obtaining anti-VWF antibodies**

Anti-VWF antibodies were obtained by immunizing human antibody-producing mice. As immunogens, keyhole limpet hemocyanin (KLH)-conjugated peptides derived from the human VWF A2 domain were used. The immunogens were administered to the mice for a total of four to six immunizations. After the final immunization, lymph nodes or spleens were collected. The lymphocytes were used to generate hybridomas by fusion with mouse myeloma cells. Culture supernatants from the hybridomas were screened by ELISA using a VWF peptide antigen. Hybridomas showing binding activity to human VWF were selected and subjected to single-cell cloning. Total RNA was extracted from the established monoclonal hybridomas, and cDNA was synthesized. Heavy-chain and light-chain antibody gene fragments were amplified by polymerase chain reaction (PCR) and sequenced. The amino acid sequences of the heavy-chain variable region and light-chain variable region of each anti-VWF antibody were thereby determined.

**SPR analysis of ADAMTS13-MDTCS binding to recombinant VWF A2**

Binding of ADAMTS13-MDTCS to recombinant VWF A2 was analyzed by SPR using HBS-EP+ buffer (Cytiva, MA, USA) as the running buffer. An anti-RFP antibody (Rockland Immunochemicals, PA, USA) was immobilized onto a CM5 sensor chip (Cytiva, MA, USA) by amine coupling using an Amine Coupling Kit (Cytiva, MA, USA). Recombinant mCherry-tagged VWF A2 was captured on the immobilized anti-RFP antibody surface before each analyte injection. mCherry-VWF A2 was diluted to 2 μg/mL in HBS-EP+ buffer and captured for 30 s. A reference flow cell without captured VWF A2 was used for background subtraction.

ADAMTS13-MDTCS was injected as the analyte using a serial dilution with a top concentration of 200 nM. Association and dissociation were monitored for 120 s and 180 s, respectively. After each cycle, the surface was regenerated with 10 mM glycine-HCl, pH 1.5 (Cytiva, MA, USA) for 240 s. Sensorgrams were reference-subtracted and analyzed for binding responses. Binding kinetics were determined using a single-cycle kinetics method, and sensorgrams were fitted using a 1:1 binding model.

**ADAMTS13-deficient mouse model**

ADAMTS13-deficient mice were used to evaluate the pharmacodynamic activity of ADAMTS13 mimetics without contribution from endogenous ADAMTS13 activity. Mice were intravenously administered ADAMTS13 mimetics or recombinant ADAMTS13 at the indicated doses, or vehicle, followed within 5 min by intravenous administration of recombinant human VWF at 400 U/kg. Twenty-four hours after the recombinant human VWF challenge, blood was collected into anticoagulant-containing tubes, and platelet counts were measured using an automated hematology analyzer.

**Supplemental Table 1. Amino acid sequences of recombinant VWF A2 substrate proteins and ADAMTS13 metalloproteinase domain construct**

| Protein | Sequence |
|---|---|
| hVWF A2-WT | HHHHHHHHENLYFQGTMVLDVAFVLEGSDKIGEADFNRSKEFMEEVIQRMDVGQDSIHVTVLQYSYMVTVEYPFSEAQSKGDILQRVREIRYQGGNRTNTGLALRYLSDHSFLVSQGDREQAPNLVYMVTGNPASDEIKRLPGDIQVVPIGVGPNANVQELERIGWPNAPILIQDFETLPREAPDLVLQR**CC** |
| hVWF A2-locked | HHHHHHHHENLYFQGT**C**SMVLDVAFVLEGSDKIGEADFNRSKEFMEEVIQRMDVGQDSIHVTVLQYSYMVTVEYPFSEAQSKGDILQRVREIRYQGGNRTNTGLALRYLSDHSFLVSQGDREQAPNLVYMVTGNPASDEIKRLPGDIQVVPIGVGPNANVQELERIGWPNAPILIQDFETLPREAPDLVLQR**C**S |
| mCherry-hVWF A2-WT | HHHHHHHHENLYFQGTMVSKGEEDNMAIIKEFMRFKVHMEGSVNGHEFEIEGEGEGRPYEGTQTAKLKVTKGGPLPFAWDILSPQFMYGSKAYVKHPADIPDYLKLSFPEGFKWERVMNFEDGGVVTVTQDSSLQDGEFIYKVKLRGTNFPSDGPVMQKKTMGWEASSERMYPEDGALKGEIKQRLKLKDGGHYDAEVKTTYKAKKPVQLPGAYNVNIKLDITSHNEDYTIVEQYERAEGRHSTGGMDELYKMVLDVAFVLEGSDKIGEADFNRSKEFMEEVIQRMDVGQDSIHVTVLQYSYMVTVEYPFSEAQSKGDILQRVREIRYQGGNRTNTGLALRYLSDHSFLVSQGDREQAPNLVYMVTGNPASDEIKRLPGDIQVVPIGVGPNANVQELERIGWPNAPILIQDFETLPREAPDLVLQR**CC** |
| mCherry-hVWF A2-locked | HHHHHHHHENLYFQGTMVSKGEEDNMAIIKEFMRFKVHMEGSVNGHEFEIEGEGEGRPYEGTQTAKLKVTKGGPLPFAWDILSPQFMYGSKAYVKHPADIPDYLKLSFPEGFKWERVMNFEDGGVVTVTQDSSLQDGEFIYKVKLRGTNFPSDGPVMQKKTMGWEASSERMYPEDGALKGEIKQRLKLKDGGHYDAEVKTTYKAKKPVQLPGAYNVNIKLDITSHNEDYTIVEQYERAEGRHSTGGMDELYK**C**SMVLDVAFVLEGSDKIGEADFNRSKEFMEEVIQRMDVGQDSIHVTVLQYSYMVTVEYPFSEAQSKGDILQRVREIRYQGGNRTNTGLALRYLSDHSFLVSQGDREQAPNLVYMVTGNPASDEIKRLPGDIQVVPIGVGPNANVQELERIGWPNAPILIQDFETLPREAPDLVLQR**C**S |
| ATS13-M-$His_6$ | AAGGILHLELLVAVGPDVFQAHQEDTERYVLTNLNIGAELLRDPSLGAQFRVHLVKMVILTEPEGAPNITANLTSSLLSVCGWSQTINPEDDTDPGHADLVLYITRFDLELPDGNRQVRGVTQLGGACSPTWSCLITEDTGFDLGVTIAHEIGHSFGLEHDGAPGSGCGPSGHVMASDGAAPRAGLAWSPCSRRQLLSLLSAGRARCVWDPPREQKLISEEDLNMHTGHHHHHH |

Amino acid sequences of recombinant hVWF A2-WT, hVWF A2-locked, mCherry-hVWF A2-WT, mCherry-hVWF A2-locked, and ATS13-M-$His_6$ are shown. The VWF A2 domain sequence is underlined. Cysteine residues introduced or retained in the A2 domain to form the intramolecular disulfide bond in A2-locked are shown in bold.

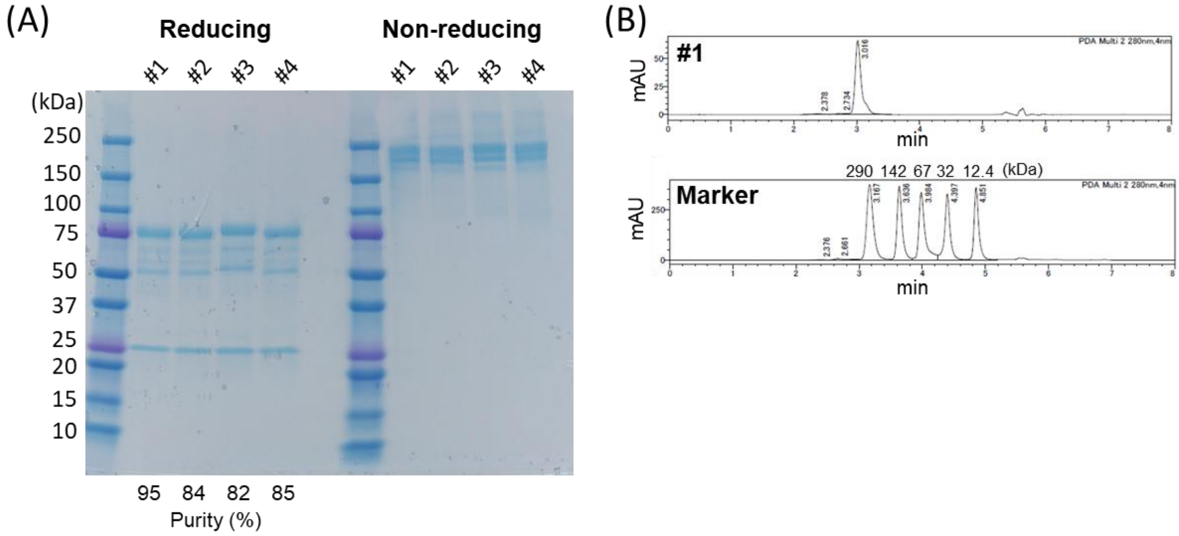


**Supplemental Figure 1. SDS-PAGE and SEC analysis of purified ADAMTS13 mimetics.** (A) Purified ADAMTS13 mimetics were analyzed by SDS-PAGE under reducing and non-reducing conditions. The molecular weight marker is shown in the indicated lanes. Protein purity was calculated from the area of the main SEC peak in panel B. (B) Representative SEC profile of ADAMTS13 mimetics with the molecular weight marker run under the same conditions. The apparent molecular size of ADAMTS13 mimetic #1 by SEC differed from its theoretical molecular weight of approximately 200 kDa, likely due to the hydrodynamic properties of the fusion protein. SEC indicates size exclusion chromatography.

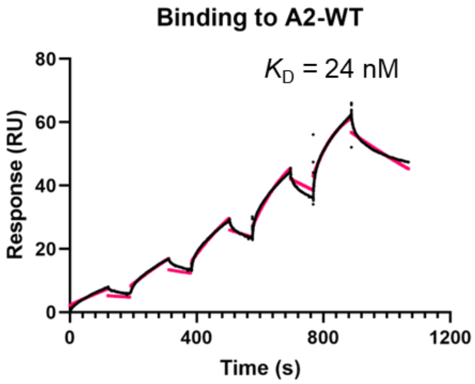


**Supplemental Figure 2. SPR analysis of ADAMTS13-MDTCS binding to recombinant VWF A2.**

Recombinant mCherry-tagged VWF A2 was captured on an anti-RFP antibody-immobilized sensor surface, and ADAMTS13-MDTCS was injected as the analyte. mCherry-tagged VWF A2 was captured at 2 μg/mL for 30 s. ADAMTS13-MDTCS was injected at a top concentration of 200 nM with a two-fold serial dilution. Association and dissociation were monitored for 120 s and 180 s, respectively. The sensor surface was regenerated with 10 mM glycine-HCl, pH 1.5, for 240 s. Representative sensorgrams are shown. Black lines indicate reference-subtracted sensorgrams, and red lines indicate fitted curves generated using a 1:1 binding model. Kinetic parameters are shown as mean values calculated from three independent biological replicates.

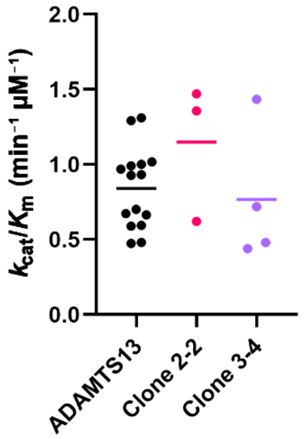


**Supplemental Figure 3. Apparent catalytic efficiency of selected ADAMTS13 mimetics measured using FRETS-VWF73.**

Apparent catalytic efficiency was evaluated using the fluorogenic FRETS-VWF73 substrate and expressed as $k_{cat}/K_m$. ADAMTS13 was included as a reference control. Clone 2-2 denotes an ADAMTS13 mimetic derived from a Peptide 2-recognizing antibody, whereas Clone 3-4 denotes ADAMTS13 mimetics derived from Peptide 3-recognizing antibodies. Each point represents one independent experiment. Horizontal bars indicate mean values. $n$ = 15 for ADAMTS13, $n$ = 3 for Clone 2-2, and $n$ = 4 for Clone 3-4.

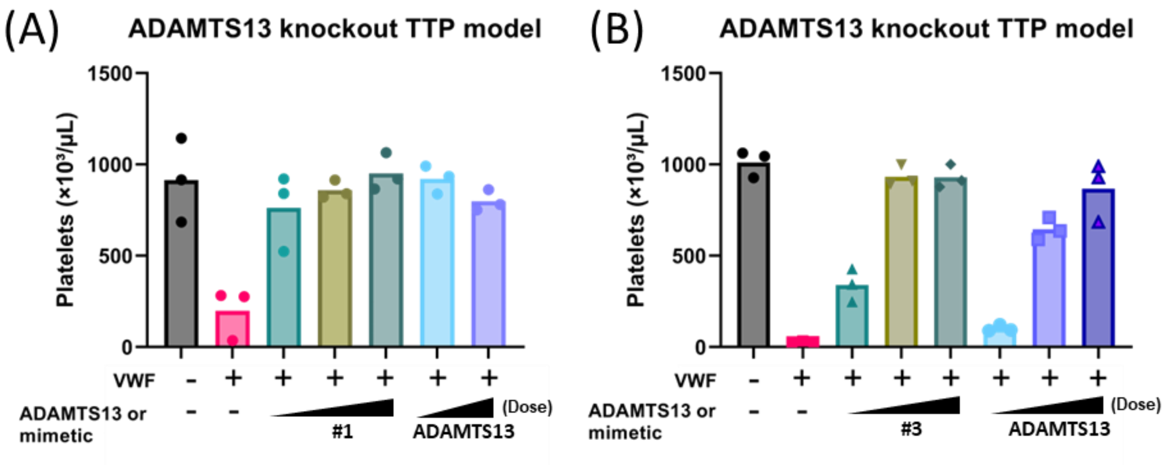


**Supplemental Figure 4. ADAMTS13 mimetics prevent VWF-induced thrombocytopenia in ADAMTS13-deficient mice.**

(A) Platelet counts in ADAMTS13-deficient mice treated with recombinant human VWF and either ADAMTS13 mimetic #1 or recombinant ADAMTS13. Mice received vehicle, ADAMTS13 mimetic #1 at 1, 5, or 10 mg/kg, or recombinant ADAMTS13 at 0.1 or 0.5 mg/kg, followed by recombinant human VWF at 400 U/kg. Platelet counts were measured 24 h after the VWF challenge. *n* = 3 mice per group. Data are shown as individual mice, with bars indicating the mean. (B) Platelet counts in ADAMTS13-deficient mice treated with recombinant human VWF and either ADAMTS13 mimetic #3 or recombinant ADAMTS13. Mice received vehicle, ADAMTS13 mimetic #3 at 0.1, 0.5, or 1 mg/kg, or recombinant ADAMTS13 at 0.01, 0.05, or 0.1 mg/kg, followed by recombinant human VWF at 400 U/kg. Platelet counts were measured 24 h after the VWF challenge. *n* = 3 mice per group. Data are shown as individual mice, with bars indicating the mean.

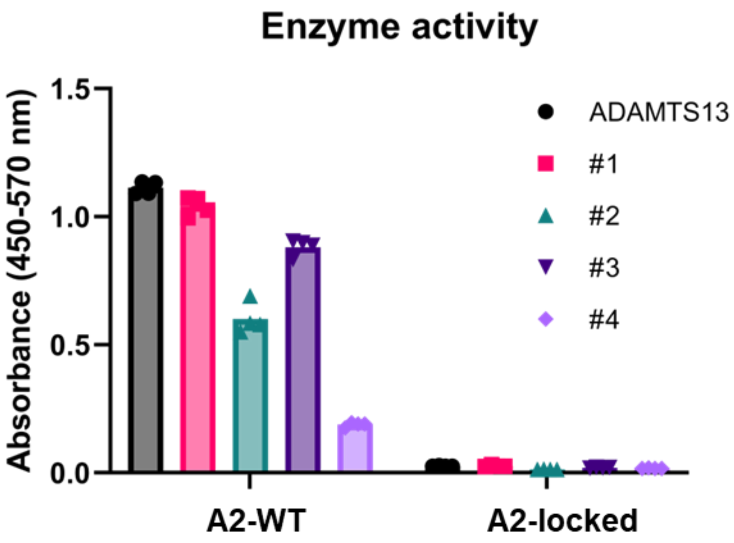


**Supplemental Figure 5. Cleavage of A2-WT and A2-locked by ADAMTS13 and ADAMTS13 mimetics.**

VWF A2-cleaving activity toward A2-WT and A2-locked was measured using the ELISA-based VWF A2 cleavage assay. Plates were coated with anti-His antibody at 5 µg/mL, and N-terminal $His_8$-tagged A2-WT or A2-locked was captured at 2 µg/mL. The immobilized VWF A2 domain was incubated with ADAMTS13 or ADAMTS13 mimetics at 100 nM for 4 h. Cleavage of the VWF A2 domain was detected using an HRP-labeled antibody recognizing the ADAMTS13-cleaved VWF A2 domain. Data are shown as mean from four biological replicates.

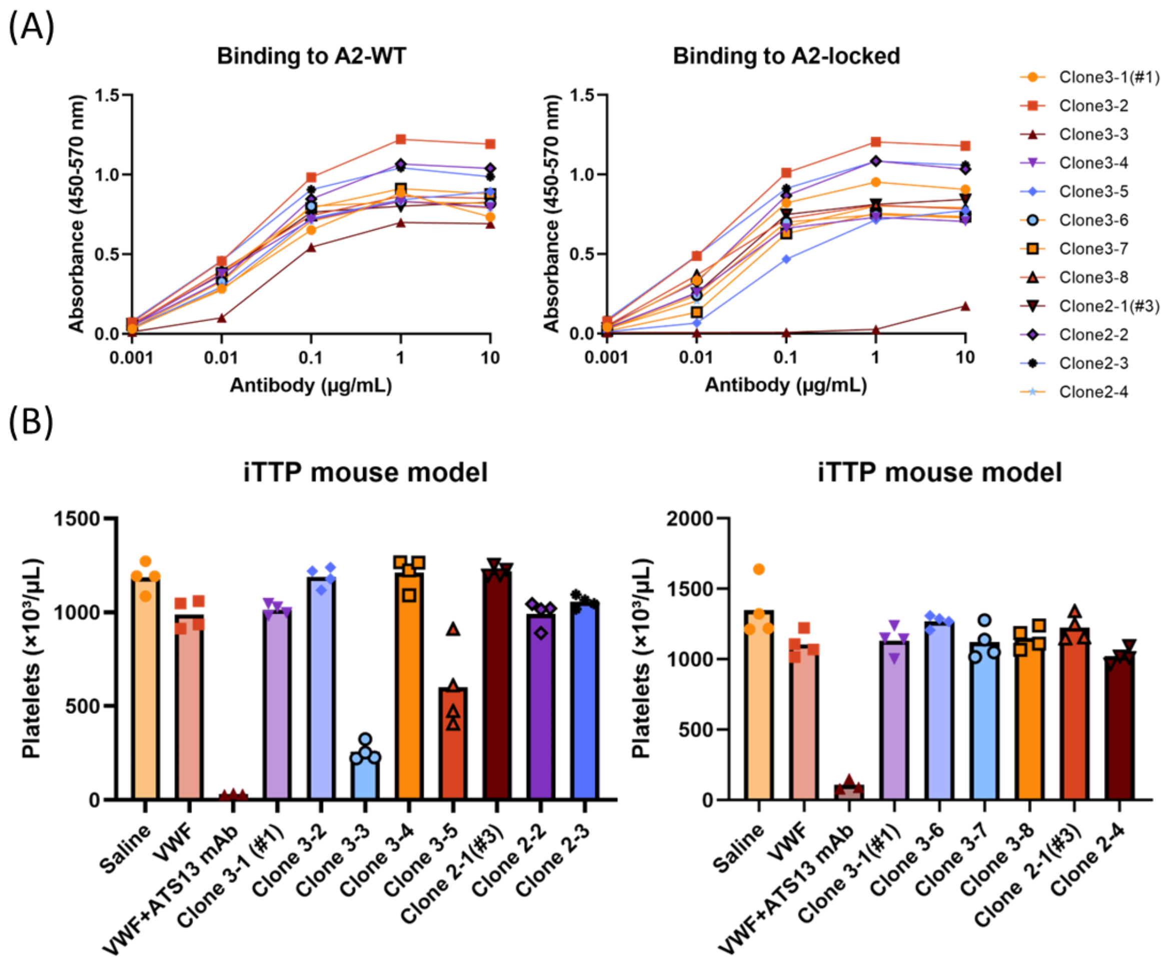


**Supplemental Figure 6. A2-locked/A2-WT binding selectivity of parental anti-VWF A2 antibodies and platelet recovery induced by the corresponding ADAMTS13 mimetics.**

(A) Binding of parental anti-VWF A2 monoclonal antibodies to A2-WT and A2-locked measured by ELISA. A2-WT or A2-locked was directly immobilized on plates at 2 µg/mL and incubated with serial dilutions of parental monoclonal antibodies. Bound antibodies were detected using an HRP-conjugated secondary antibody. Data are shown as mean values from two independent experiments. (B) Platelet recovery induced by the corresponding ADAMTS13 mimetics in the iTTP mouse model after the recombinant human VWF challenge. Mice in the ADAMTS13 mimetic-treated groups received the inhibitory anti-ADAMTS13 antibody mixture followed by the recombinant human VWF challenge. The inhibitory anti-ADAMTS13 antibody mixture was administered at a total dose of 30 mg/kg, recombinant human VWF at 500 U/kg, and ADAMTS13 mimetics at 5 mg/kg. Data are shown as mean values from four mice per group.
